# Antibacterial activity of tamoxifen derivatives against methicillin-resistant *Staphylococcus aureus*

**DOI:** 10.1101/2024.07.16.603795

**Authors:** Irene Molina Panadero, Javier Falcón Torres, Karim Hmadcha, Salvatore Princiotto, Luigi Cutarella, Mattia Mori, Sabrina Dallavalle, Michael S. Christodoulou, Younes Smani

## Abstract

The present work aimed to discover new tamoxifen derivatives with antimicrobial potential, particularly targeting methicillin-resistant *Staphylococcus aureus* (MRSA).

The MIC of 22 tamoxifen derivatives was determined against *S. aureus* reference and MRSA strains, using microdilution assays. The antibacterial effects of selected tamoxifen derivatives against MRSA (USA7) were assessed through bacterial growth assays. Bacterial membrane permeability and molecular docking assays were performed.

The MIC of the tamoxifen derivatives against MRSA ranged from to 16 to >64 μg/mL. Bacterial growth assays demonstrated that tamoxifen derivatives **2**, **5**, and **6** reduced dose-dependently the growth of the USA7 strain. Moreover, treatment of MRSA with derivatives **2** and **5** resulted in increased membrane permeabilization without being the cell wall their molecular target.

These data suggest that tamoxifen derivatives exhibit antibacterial activity against MRSA, potentially broadening the spectrum of available drug treatments for combating antimicrobial-resistant Gram-positive bacteria.

**Importance:** The development of new antimicrobial therapeutic strategies requires immediate attention to avoid the tens of millions of deaths predicted to occur by 2050 as a result of multidrug-resistant (MDR) bacterial infections. In this study, we assessed the antibacterial activity of 22 tamoxifen derivatives against methicillin-resistant Staphylococcus aureus (MRSA). We found that three tamoxifen derivatives exhibited antibacterial activity against MRSA clinical isolats, presenting MIC_50_ values between 16 and 64 μg/mL and reducing bacterial growth over 24 h. Additionally, this antibacterial activity for two of the derivatives was accompanied by increased membrane permeability of MRSA. Our results suggest that tamoxifen derivatives might be used as a potential therapeutic alternative for treating MRSA strains in an animal model of infection.

## INTRODUCTION

In the last decades, Gram-positive bacteria have demonstrated an increasing antimicrobial resistance, driven by various genomic, transcriptomic, and proteomic adaptations^1^. This alarming trend aligns with a concerning decline in the development of new antibiotics, a phenomenon often referred to as the “Post-Antibiotic Era”^2^. This situation underscores the urgent need for effective solutions, a concern highlighted by numerous institutions (ref). Consequently, there is a growing demand for innovative antimicrobial therapeutic approaches, including the exploration of non-antibiotic compounds and drug repurposing, both as monotherapy and in combination with the limited clinically relevant antibiotics currently available^3^.

Among the various strategies employed in the discovery of new antibacterial agents, drug repurposing is one of the most exploited^3^. A very representative example of such approach is tamoxifen, a well-known chemotherapic drug, widely used for decades as the gold standard for the treatment of estrogen receptor positive breast cancers and related metastatic forms^4^. Recently, tamoxifen has exhibited relevant antibacterial properties against a range of pathogenic microorganisms, including Gram-positive *Staphylococcus epidermidis*, and *Enterococcus faecalis* and Gram-negative *Escherichia coli* and *Acinetobacter baumannii*^5,6^. This antimicrobial effect may be ascribed to the cytochrome P450-mediated metabolism of tamoxifen, resulting in the generation of three major metabolites: *N*-desmethyltamoxifen, 4-hydroxytamoxifen and endoxifen^5,7^. Only a limited number of studies have reported the activity of these metabolites against various infectious agents^5,6,8–12^. Among these, 4-hydroxytamoxifen has garnered attention for its chemical behavior as a weak base, since it has been observed that such property is responsible for the protection of cells and mice against lethal Shiga toxin 1 (STx1) or Shiga toxin 2 (STx2) toxicosis^8^. It has also demonstrated efficacy against *Plasmodium falciparum* and *Cryptococcus neoformans*^10,11^. Furthermore, when used in monotherapy, 4-hydroxytamoxifen has displayed activity against *Mycobacterium tuberculosis* (with a MIC_50_ of approximately 2.5 to 5 μg/mL)^12^. Endoxifen’s activity was studied against *C. neoformans*, revealing a MIC of 4 μg/mL^11^. The combination of *N*-desmethyltamoxifen, 4-hydroxytamoxifen and endoxifen exhibited MIC_50_ values of 8 and 16 μg/mL against clinical isolates of *A. baumannii* and *E. coli*, and MIC_50_ values of 1 and 2 μg/mL against clinical isolates of *S. epidermidis* and *E. faecalis*^5,6^.

To identify new chemical compounds with antimicrobial potential, a collection of 22 tamoxifen derivatives, bearing different electron-withdrawing or electron-donating substituents on the aromatic rings A and B (Table 1) was tested against methicillin-resistant *S. aureus* (MRSA).

**Table 1.** MIC (μg/mL) of tamoxifen derivatives **1** – **22** for reference strain of *S. aureus*

| Compound | Structure | R<sub>1</sub> | R<sub>2</sub> | R<sub>3</sub> | R<sub>4</sub> | *S. aureus*<br>ATCC 1556 |
| --- | --- | --- | --- | --- | --- | --- |
| **1** | A | H | H | H | H | >64 |
| **2** | A | OH | CON(CH<sub>2</sub>CH<sub>3</sub>)<sub>2</sub> | H | OH | **32** |
| **3** | A | OBn | COCH<sub>2</sub>CH<sub>3</sub> | H | OBn | >64 |
| **4** | B | OBn | COCH<sub>2</sub>CH<sub>3</sub> | H | OBn | >64 |
| **5** | A | OH | CON(CH<sub>3</sub>)<sub>2</sub> | H | OH | **32** |
| **6** | B | OH | CON(CH<sub>3</sub>)<sub>2</sub> | H | OH | **16** |
| **7** | A | OCH<sub>3</sub> | CO*i*Propyl | H | H | >64 |
| **8** | B | OCH<sub>3</sub> | CO*i*Propyl | H | H | >64 |
| **9** | A | H | CO*i*Propyl | H | H | >64 |
| **10** | B | H | CO*i*Propyl | H | H | >64 |
| **11** | B | H | CO*i*Propyl | H | OBn | >64 |
| **12** | A | H | CO*i*Propyl | H | OBn | >64 |
| **13** | A | OH | CO*i*Propyl | Cl | Cl | >64 |
| **14** | B | OH | CO*i*Propyl | Cl | Cl | >64 |
| **15** | A | OCH<sub>3</sub> | CO*i*Propyl | H | OBn | >64 |
| **16** | B | OCH<sub>3</sub> | CO*i*Propyl | H | OBn | >64 |
| **17** | B | OCH<sub>3</sub> | CO*i*Propyl | H | OH | >64 |
| **18** | A | OCH<sub>3</sub> | CO*i*Propyl | H | OH | >64 |
| **19** | A | OH | CO*i*Propyl | H | H | >64 |
| **20** | B | OH | CO*i*Propyl | H | H | >64 |
| **21** | A | OBn | CO*i*Propyl | H | OBn | >64 |
| **22** | B | OBn | CO*i*Propyl | H | OBn | >64 |

## RESULTS

### Antimicrobial activity of tamoxifen derivatives

Twenty-two tamoxifen derivatives (Table 1) were subjected to evaluation to determine their MIC against *S. aureus* ATCC 1556 reference strain. Only the tamoxifen derivatives **2**, **5** and **6** showed MIC for *S. aureus* ATCC 1556 strain ranging from 16 to 32 μg/mL, while the rest of the compounds did not show MIC values <64 μg/mL (Table 1).

A comparison between the biological results and the chemical structures of the tamoxifen derivatives allowed some interesting structure-activity relationships outcomes. In particular, the most active compounds **2**, **5** and **6** share the presence of the hydroxyl group in *para* position on both phenyl rings, A and B. All the derivatives bearing the hydroxyl group on only one of the phenyl rings resulted inactive (compounds **13**, **14**, **17**, **18**, **19** and **20**). In the case of no substituents on the phenyl rings (compounds **1**, **9** and **10**) or in presence of electron-donating substituents diverse from the hydroxyl group on one or both phenyl rings A and B, the derivatives resulted inactive as well (compounds **3**, **4**, **7**, **8**, **11**, **12**, **15**, **16**, **21** and **22**).

The most promising derivatives were selected on the base of a MIC ≤64 μg/mL and were evaluated against 7 clinical isolates of MRSA to extend their biological evaluation to the antimicrobial effect on resistant strains. As shown in Table 2, the MICs of tamoxifen derivatives **2**, **5** and **6** for MRSA strains ranged from 16 to 64 μg/mL.

**Table 2.**
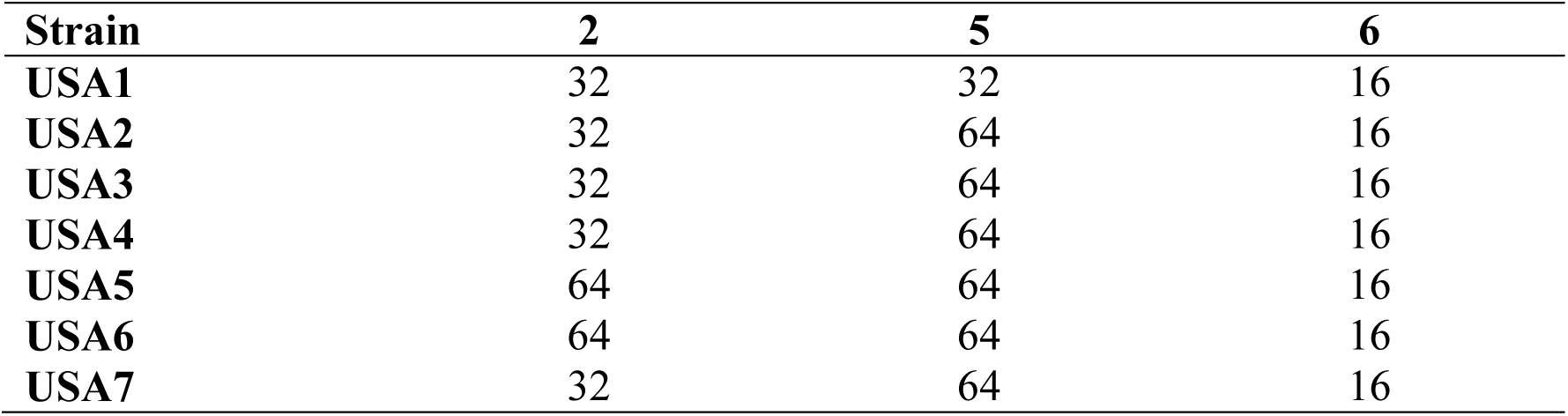
MIC of tamoxifen derivatives **2**, **5** and **6** for clinical isolates for clinical methicillin-resistant *S. aureus*.

### Bacterial growth curves

Using bacterial growth, we examined the antibacterial activity of the selected tamoxifen derivatives against MRSA USA7 strain (Figure 1). Compounds **2**, **5** and **6** at concentrations of 1x, 2x and 4x MIC, reduced the growth of this strain in a concentration-dependent manner during 24 h. This antibacterial effect is more pronounced at 2x and 4xMIC.

**Figure 1.**
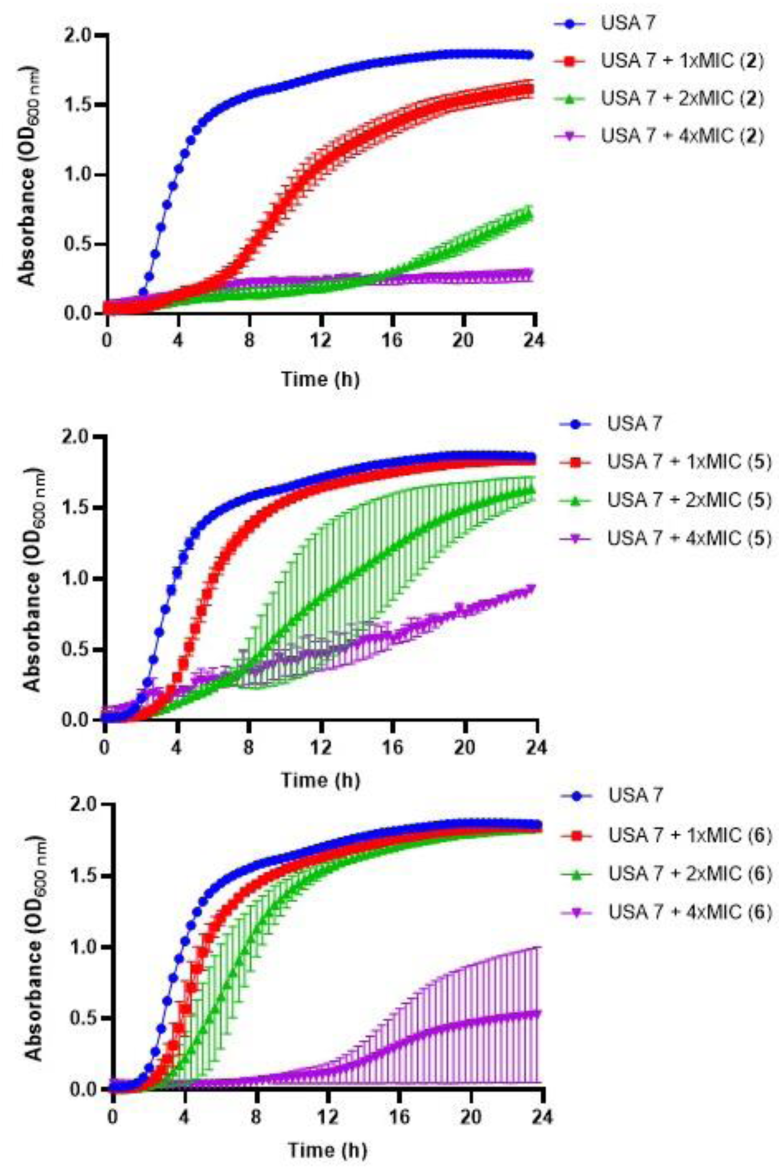
Antibacterial activity of tamoxifen derivatives against clinical methicillin-resistant *S. aureus*. Bacterial growth curves of MRSA USA7 strain in the presence of 1x, 2x and 4xMIC of tamoxifen derivative **2, 5 or 6** for 24 h. Data are represented as mean from two independent. experiments. COL: colistin.

### Effect of tamoxifen derivatives on the bacterial cell membrane

In order to determine the mode of action of the selected tamoxifen derivatives, their effect on the membrane permeability of USA7 strain was evaluated by incubation with ethidium homodimer 1, a fluorescent marker known to enter bacterial cells when the membrane integrity is compromised.

Fluorescence monitoring using a Typhoon FLA scanner for 3 h showed a slight increase in the cellular fluorescence of the USA7 strain when treated with sub-MIC of the tamoxifen derivatives **2** and **5**, but not for **6**. (Figure 2).

**Figure 2.**
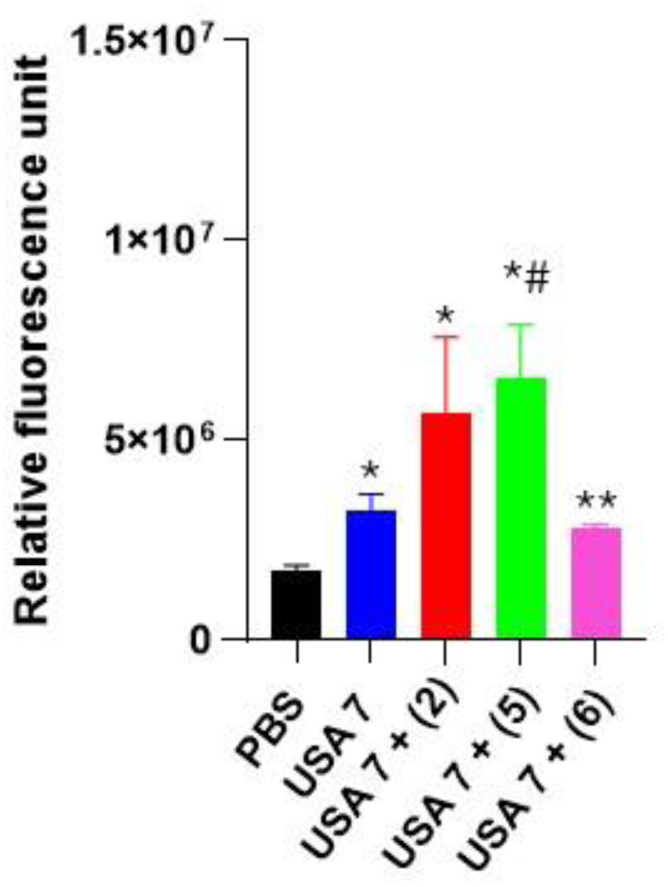
Effect of tamoxifen derivatives on the bacterial permeability against clinical methicillin-resistant *S. aureus.* The membrane permeabilization of MRSA USA7 strain in the presence of 1xMIC of tamoxifen derivative **2, 5** or **6**, incubated for 10 min, was quantified by Typhon Scanner. Data are represented as mean ± SEM from three independent experiments. \**P* < 0.05 vs PBS, #*P* < 0.05 vs USA7, \**P* < 0.05 vs USA7 + (5).

### Tamoxifen derivatives interaction with bacterial membrane

To investigate the interaction between tamoxifen derivates **5** and **6** and the bacterial membrane model, and to assess how subtle structural modifications (i.e., *cis*/*trans* conformation) might affect their ability to embed within the target, unbiased molecular dynamics (MD) simulations were carried out starting from the small molecules being placed in the solvent area.^13–19^

Different from previous findings with different scaffolds^17^, for derivatives **5** and **6** the density plots clearly evidence that the molecules are unable to penetrate deeply into the membrane model (Figure 3A,B). In fact, density peaks of the two derivatives slightly exceeds the density peak of the phosphates that represent the membrane outermost layer. These results suggest that the *S. aureus* membrane might not be the molecular target of **5** and **6** antibacterial efficacy.

**Figure 3:**
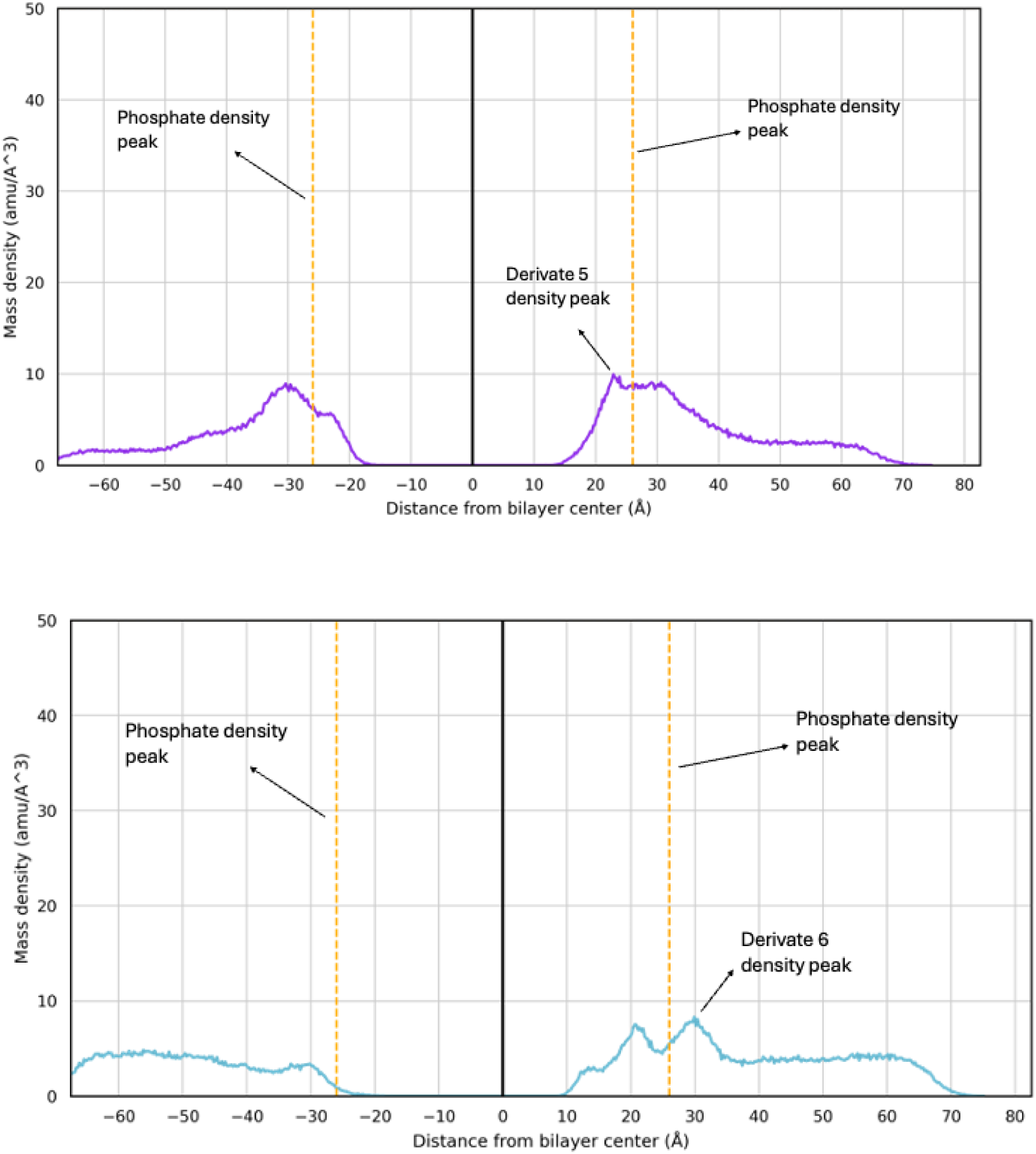
Mass density profile of **5 (A)** and **6 (B)** with the corresponding density peaks highlighted. The density peaks of the lipid phosphate groups are shown as orange dashed lines, with maximum density at −26 Å and +26 Å.

To further provide atomistic details of this outcome, the distance between each molecule and the lipid bilayer was monitored along MD trajectories. Then, the MD frame corresponding to the smallest intermolecular distance was extracted and visually inspected. A labile H-bond interaction between the phenol moiety of the molecules and the protonated ammino group (i.e., R-NH_3_^+^) of a phosphatidylglycerol residues at the membrane interface with the solvent was observed (Figure 4A,B). Unlike **5**, derivative **6** also establishes an additional interaction with the -OH group of a phosphatidylglycerol residue (Figure 4B). However, this interaction is not stable in time and failed to promote molecules embedding into the *S. aureus* membrane model within the simulation time, which further reinforce the hypothesis that **5** and **6** exploit their antibacterial potentials through the interaction with target that differ from the bacterial membrane.

**Figure 4:**
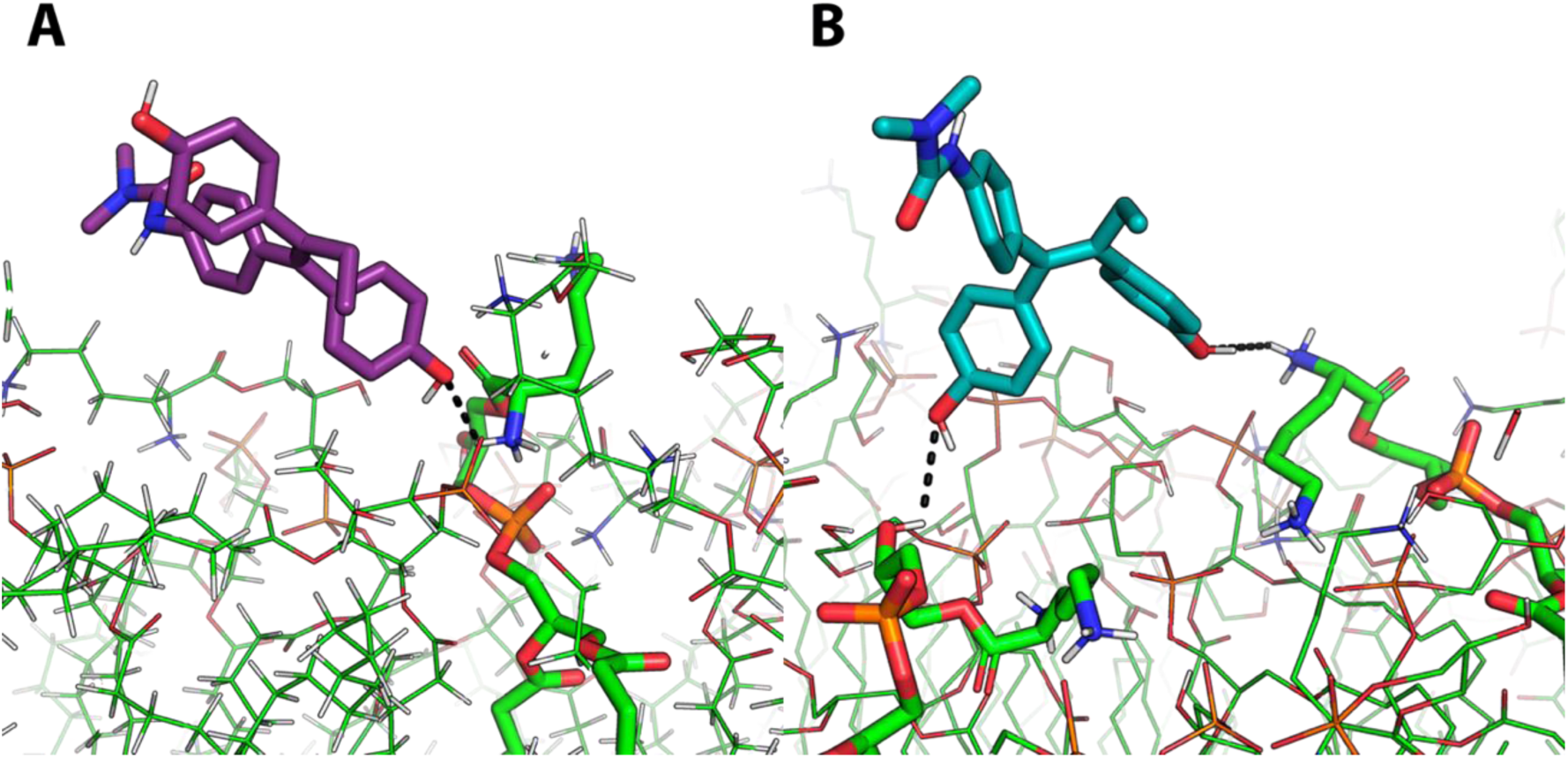
Representative MD frames corresponding to the minimum distances between derivate **5** (**A**, purple sticks) or derivate **6** (**B**, light blue sticks) and the *S. aureus*membrane model. The interactions between these two derivates and the NH3^+^ moiety (panel **A** and **B**) and -OH moiety (panel **B**) of phosphatidylglycerol residues is represented by black dashed lines.

## DISCUSSION

The rise of multidrug resistant *S. aureus* has led to a wide use of the last options for the treatment of severe infections caused by this microorganism, and to the consequent acquired antimicrobial resistance worldwide in the last decades^20^. In previous studies, tamoxifen and its metabolites garnered attention as a potential repurposed drug for treatment of infectious diseases^9^.

Due to the antibacterial effect of tamoxifen and its metabolites *N*-desmethyltamoxifen, 4-hydroxytamoxifen and endoxifen against *S. epidermidis* and *A. baumannii*^5,6^, we hypothesized that tamoxifen derivatives may enhance this antibacterial activity against MRSA. For this purpose, twenty-two tamoxifen derivatives were tested against *S. aureus* reference strain.

Three tamoxifen derivatives **2**, **5** and **6** in monotherapy showed antibacterial activity against MRSA strains being the MIC between 16 and 64 μg/mL, which overall fell within the range of other known antibiotics such as amikacin, teicoplanin, sulfonamides nitrofurantoin^21,22^. However, none of the rest of tested tamoxifen derivatives exhibited activity against MRSA. Reasons for this difference could be related to the chemical structure of these derivatives, being **2**, **5** and **6** the only compounds bearing the electron-donating hydroxyl group in *para* position on both phenyl rings A and B.

The antibacterial activity of tamoxifen derivatives **2** and **5** at 1x, 2x and 4x MIC against MRSA USA7 strain was more pronounced and began earlier than with tamoxifen derivative **6**. This result may be related to the *trans* stereochemistry (structure **A**) of compounds **2** and **5** and their ability to act on the cell wall of *S. aureus*, as observed with tamoxifen metabolites against Gram-negative bacteria^5^. Permeability assays confirm this observation, since tamoxifen derivatives **2** and **5**, but not **6**, produced a slight increase in membrane permeability of MRSA, suggesting that the bacterial cell wall integrity could be affected and could acts as the target of both derivatives. In support of this hypothesis, tamoxifen has been shown to increase the membrane permeability of *Streptococcus pneumoniae* by perturbing the phospholipid bilayer^23^.

Even though tamoxifen derivatives might affect cell wall permeability, MD simulations have reported that the *S. aureus* membrane might not be the molecular target of these derivatives due to their instability over time. Alternative mechanisms of action, rather than the perturbation of the cell membrane, cannot be ruled out. Of note, in other microorganisms such as fungi, the mechanism of action of the tamoxifen is well-documented and involves its binding to calmodulin^24,25^. Moreover, the tamoxifen metabolite, 4-hydroxytamoxifen, can potentially inhibit bacterial phospholipase D in *P. aeruginosa*^26^. Additional research focusing on understanding the mechanism of action of the tested tamoxifen derivatives against MRSA would be of significant interest since different chemical structures resulted active on the aforementioned strain, thus providing a better therapeutic efficacy.

The antimicrobial activity of the selected derivatives identified in this work hints at a promising potential that deserves to be further explored *in vivo* after determining their pharmacokinetic parameters. However, *in vitro* bacterial growth showed a progressive regrowth of the MRSA USA 7 strain after treatment with these tamoxifen derivatives, suggesting that acquired resistance could take place. Nevertheless, it should be noted that the MICs of tamoxifen derivatives **2**, **5**, and **6** against the USA 7 strain in this bacterial growth condition are 32, 32, and 16 μg/mL, respectively, which are below the original 2x and 4x MIC of the tamoxifen derivatives concentration. Further investigations, including the determination of tamoxifen derivatives concentration during the bacterial growth assay, are necessary to better understand the regrowth of this strain in their presence.

In conclusion, this study’s findings offer fresh perspectives on the application of a diverse group of tamoxifen derivatives against MRSA, pathogen for which treatment options are dramatically restricted. Despite the promising potential displayed by this novel class of antibacterial compounds, additional research is required to uncover the mechanism of action of these chemical entities. Moreover, determining the ideal dosage that ensures therapeutic effectiveness in managing severe infections caused by multidrug-resistant Gram-positive remains a critical objective.

## MATERIAL AND METHODS

### Bacterial Strains

A total of 7 MRSA clinical isolates and one *S. aureus* reference ATCC 1556 strain were used in this study^27^.

### Antimicrobial agent and tamoxifen derivatives

The small library of tamoxifen derivatives was prepared as described in literature, starting from appropriately substituted benzophenones which underwent McMurry olefination^28,29^. The structures of the tested compounds are presented in Table 1.

### *In vitro* susceptibility testing

The MICs of tamoxifen derivatives were determined against reference and MRSA strains in two independent experiments using the broth microdilution method, in accordance with the standard guidelines of the European Committee on Antimicrobial Susceptibility Testing (EUCAST)^30^. A 5x10^5^ cfu/mL inoculum of each strain was cultured in Luria Bertani (LB) and added to U bottom microtiter plates (Deltlab, Spain) containing the studied tamoxifen derivatives. The plates were incubated for 18 h at 37 °C. *Pseudomonas aeruginosa* ATCC 27853 was used as the positive control strain.

### Bacterial growth curves

To determine the antibacterial and synergistic effects, duplicate bacterial growth curves were performed for MRSA USA7 strain. A 1/200 dilution of an overnight bacterial cultures grown in LB at 37 °C with continuous agitation at 180 rpm was performed in LB in 96-well plate in the presence of 1x, 2x and 4x MIC of compounds **2**, **5** and **6**. A drug-free broth was evaluated in parallel as control. Absorbance measurements at 600 nm every 20 min for 24 h were conducted using a Tecan spectrophotometer (model XYZ-2000, Austria).

### Membrane permeability assays

The bacterial cells were grown in LB broth and incubated in the absence or presence of (i) 1x MIC of compounds **2**, **5** and **6** for 3 h. The pellet was harvested by ultracentrifugation at 4600g for 15 min. The bacterial cells were washed with PBS 1X, and after centrifugation in the same condition described before, the pellet was resuspended in 100 µL of PBS 1X containing 10 μL of Ethidium Homodimer-1 (ThermoFisher, USA). After 10 min of incubation, 100 μL was placed into a 96-well plate to measure fluorescence for 300 min using a Typhoon FLA 9000 laser scanner (GE Healthcare Life Sciences, USA) and quantified using ImageQuant TL software (GE Healthcare Life Sciences, USA)^31^.

### Molecular docking assays

The symmetric lipid bilayer membrane of *S. aureus* was built using the CHARMM-GUI Membrane Builder Tool^13,14^. This system was designed to mimic the phospholipid composition of the *S. aureus* membrane, which consists of 56.8% phosphatidylglycerol (PG), 37.9% Lys-PG, and 5.3% diphosphatidylglycerol (DPG), also referred as Cardiolipin (CL), in agreement with previous molecular dynamics (MD) studies^15–17^. Each lipid bilayer contained a total of 95 lipid molecules, positioned with their centers at z = 0., surrounded by a water layer with a thickness of 50 Å. The system was neutralized using Na^+^ and Cl^-^ ions at a concentration of 0.145 M, as suggested by CHARMM-GUI.

Derivatives **5** and **6** were drawn using the Sketchpad powered by Marvin JS included in the ligand reader & modeler module of CHARMM-GUI, and were parameterized using the standard CHARMM force field (FF)^32^. The Multicomponent Assembler tool was used to randomly distribute the small molecules under investigation within the solvent area. The membrane systems containing these small molecules were then parameterized with a membrane thickness of 50 Å and a box XY length of 79.50 Å^33,34^. Following system construction with CHARMM-GUI, the topology and coordinate files were generated for AMBER^35,36^.

To investigate the interaction between **5** and **6** with the *S. aureus* membrane model, MD simulations were conducted using AMBER22^37,38^. The initial system was energy minimized for a total of 40000 steps., with the first 1500 steps employing the steepest descent algorithm, followed by the conjugate gradient algorithm for the remaining steps. A non-bonded cut-off of 10Å was used. After energy minimization, each system was gradually heated to 300 K over 900 ps at constant volume using the Langevin thermostat with a collision frequency of 2 ps^-1^, and then left at 300 K at constant volume for 200 ps. Box density was equilibrated at constant pressure and constant temperature (300 K) over 1 ns using the Berendsen barostat.

Following density equilibration, a preliminary 50 ns MD simulation was conducted at constant pressure. Subsequently, MD trajectories were generated for 500 ns. In all MD simulations, no positional restraints were applied.

Analysis of MD trajectories was performed using the CPPTRAJ^39^ program from the AmberTools package. This analysis included calculation of the mass densities of derivatives within the system along the z-axis. Small molecules interactions with the membrane were visually inspected with PyMol^40^.

### Statistical Analysis

Group data are presented as means ± standard errors of the means (SEM). The student *t*-test was used to determine differences between means using the GraphPad Prism 9. A *p*-value <0.05 was considered significant.

## DATA AVAILABILITY

The data that support the findings of this study are available from the corresponding author upon reasonable request.

## ACKNOWLEDGMENTS

We thank José María Marimon Ortiz de Zárate for sharing the MRSA clinical isolates, Cayetana Martín and the Proteomic facility of the Andalusian Center of Developmental Biology for their technical help and the COST Action CA21145 – European Network for diagnosis and treatment of antibiotic-resistant bacterial infections (EURESTOP). This work was funded by the Consejería de Universidad, Investigación e Innovación de la Junta de Andalucía (grant ProyExcel_00116).

## AUTHOR CONTRIBUTIONS

Conceptualization, M.S.C. and Y.S.; methodology, I.M.P., J.F.T., M.S.C., L.C. and S.P.; formal analysis, I.M.P., J.F.T., K.H., M.M. and Y.S.; writing—original draft preparation, I.M.P. and Y.S.; writing-review and editing, M.S.C., M.M., S.D. and Y.S.; funding acquisition, Y.S. All authors have read and agreed to the published version of the manuscript.

## REFERENCES

1. Yelin, I. & Kishony, R. Antibiotic resistance. Cell 172, 1136–1136 (2018).

2. Bagley, N. & Outterson, K. We will miss antibiotics when they’re gone. The New York Times, January 18 (2017).

3. Canturri, A. M. & Smani, Y. Anthelmintic drugs for repurposing against Gram-negative bacilli infections. Curr. Med. Chem. 30, 59–71 (2022).

4. Green, K. A. & Carroll, J. S. Oestrogen-receptor-mediated transcription and the influence of co-factors and chromatin state. Nat. Rev. Cancer 7, 713–22 (2007).

5. Miró-Canturri, A. et al. Repurposing of the tamoxifen metabolites to combat infections by multidrug-resistant gram-negative bacilli. Antibiotics 10, 336 (2021a).

6. Miró-Canturri, A. et al. Repurposing of the tamoxifen metabolites to treat methicillin-resistant *Staphylococcus epidermidis* and vancomycin-resistant *Enterococcus faecalis* infections. Microbiol. Spectr. 9, e0040321 (2021b).

7. Klein, D. J. et al. PharmGKB summary: tamoxifen pathway, pharmacokinetics. Pharmacogenet. Genomics 23, 643–647 (2013).

8. Selyunin, A. S., Hutchens, S., McHardy, S. F. & Mukhopadhyay, S. Tamoxifen blocks retrograde trafficking of Shiga toxin 1 and 2 and protects against lethal toxicosis. Life Sci. Alliance 2, e201900439 (2019).

9. Montoya, M. C. & Krysan, D. J. Repurposing estrogen receptor antagonists for the treatment of infectious disease. mBio 9, e02272–18 (2018).

10. Weinstock, A. et al. Tamoxifen activity against *Plasmodium* in vitro and in mice. Malar. J. 18, 378 (2019).

11. Butts, A. et al. Estrogen receptor antagonists are anti-cryptococcal agents that directly bind EF hand proteins and synergize with fluconazole *in vivo*. mBio 5, e00765–13 (2014).

12. Chen, F. C. et al. Pros and cons of the tuberculosis drugome approach-an empirical analysis. PLoS One 9, e100829 (2014).

13. Li Y, Liu J, Gumbart JC. Preparing Membrane Proteins for Simulation Using CHARMM-GUI. In: Methods in Molecular Biology. Vol 2302; 2021.

14. Jo S, Kim T, Iyer VG, Im W. CHARMM-GUI: A web-based graphical user interface for CHARMM. J Comput Chem. 29(11) (2008).

15. Piggot TJ, Holdbrook DA, Khalid S. Electroporation of the E. coli and S. aureus membranes: Molecular dynamics simulations of complex bacterial membranes. J Physic Chemi B. 115(45) (2011).

16. Kim W, Zhu W, Hendricks GL, et al. A new class of synthetic retinoid antibiotics effective against bacterial persisters. Nature. 2018;556(7699). doi:10.1038/nature26157.

17. Princiotto S, Casciaro B, G. Temprano A, et al. The antimicrobial potential of adarotene derivatives against Staphylococcus aureus strains. Bioorg Chem. 2024;145. doi:10.1016/j.bioorg.2024.107227.

18. Witzke S, Petersen M, Carpenter TS, Khalid S. Molecular Dynamics Simulations Reveal the Conformational Flexibility of Lipid II and Its Loose Association with the Defensin Plectasin in the Staphylococcus aureus Membrane. Biochemistry. 2016;55(23). doi:10.1021/acs.biochem.5b01315.

19. Joodaki F, Martin LM, Greenfield ML. Generation and Computational Characterization of a Complex Staphylococcus aureus Lipid Bilayer. Langmuir. 2022;38(31). doi:10.1021/acs.langmuir.2c00483.

20. Abebe AA, Birhanu AG. Methicillin resistant *Staphylococcus aureus*: Molecular mechanisms underlying drug resistance development and novel strategies to combat. Infect. Drug Resist. 16, 7641–7662 (2023).

21. European Committee on Antimicrobial Susceptibility Testing. European antimicrobial breakpoints. Basel: EUCAST, https://www.eucast.org/fileadmin/src/media/PDFs/EUCAST_files/Breakpoint_tables/v_14.0_Breakpoint_Tables.pdf (2024).

22. Clinical and Laboratory Standards Institute (CLSI). Performance Standards for Antimicrobial Susceptibility Testing. 34th ed. CLSI supplement M100 (2024).

23. Ortiz-Miravalles L, Sánchez-Angulo M, Sanz JM, Maestro B. Drug repositioning as a therapeutic strategy against Streptococcus pneumoniae: Cell membrane as potential target. Int. J. Mol. Sci. 24(6):5831.

24. Dolan, K. et al. Antifungal activity of tamoxifen: in vitro and in vivo activities and mechanistic characterization. Antimicrob Agents Chemother. 53, 3337–3346 (2009).

25. Butts, A. et al. Structure-activity relationships for the antifungal activity of selective estrogen receptor antagonists related to tamoxifen. PLoS One 10, e0125927 (2015).

26. Scott, S. A. et al. Discovery of desketoraloxifene analogues as inhibitors of mammalian, *Pseudomonas aeruginosa*, and nape phospholipase D enzymes. ACS Chem Biol 10, 421–432 (2015).

27. Docobo Pérez, F. Tratamiento de la neumonía experimental por Staphylococus aureus. Estudios de eficacia terapéutica de cotrimoxazol, cloxacilina, linezolid y vancomicina frente a cepas S. aureus sensible y resistente a meticilin. Ph.D. Thesis, Seville University, Seville, Spain, 26 March 2009.

28. Christodoulou, M. S. et al. Synthesis and biological evaluation of novel tamoxifen analogues, Bioorg. Med. Chem. 21, 4120–4131 (2013).

29. Christodoulou, M. S. et al. 4-(1,2-diarylbut-1-en-1-yl)isobutyranilide derivatives as inhibitors of topoisomerase II. Eur. J. Med. Chem. 118, 79–89 (2016).

30. European Committee on Antimicrobial Susceptibility Testing European antimicrobial breakpoints. Basel: EUCAST, https://www.eucast.org/ast_of_bacteria/mic_determination (2022).

31. Miró-Canturri, A., Ayerbe-Algaba, R., Villodres, Á. R., Pachón, J. & Smani, Y. Repositioning rafoxanide to treat Gram-negative bacilli infections. J. Antimicrob. Chemother. 75, 1895–1905 (2020).

32. Kim S, Lee J, Jo S, Brooks CL, Lee HS, Im W. CHARMM-GUI ligand reader and modeler for CHARMM force field generation of small molecules. J Comput Chem. 2017;38(21). doi:10.1002/jcc.24829.

33. Huang J, Rauscher S, Nawrocki G, et al. CHARMM36m: An improved force field for folded and intrinsically disordered proteins. Nat Methods. 2016;14(1). doi:10.1038/nmeth.4067.

34. Lee J, Hitzenberger M, Rieger M, Kern NR, Zacharias M, Im W. CHARMM-GUI supports the Amber force fields. Journal of Chemical Physics. 2020;153(3). doi:10.1063/5.0012280.

35. Lee J, Cheng X, Swails JM, et al. CHARMM-GUI Input Generator for NAMD, GROMACS, AMBER, OpenMM, and CHARMM/OpenMM Simulations Using the CHARMM36 Additive Force Field. J Chem Theory Comput. 2016;12(1). doi:10.1021/acs.jctc.5b00935.

36. Brooks BR, Brooks CL, Mackerell AD, et al. CHARMM: The biomolecular simulation program. J Comput Chem. 2009;30(10). doi:10.1002/jcc.21287.

37. Salomon-Ferrer R, Case DA, Walker RC. An overview of the Amber biomolecular simulation package. Wiley Interdiscip Rev Comput Mol Sci. 2013;3(2). doi:10.1002/wcms.1121.

38. Case DA, Cheatham TE, Darden T, et al. The Amber biomolecular simulation programs. J Comput Chem. 2005;26(16). doi:10.1002/jcc.20290.

39. Roe DR, Cheatham TE. PTRAJ and CPPTRAJ: Software for processing and analysis of molecular dynamics trajectory data. J Chem Theory Comput. 2013;9(7). doi:10.1021/ct400341p.

40. Delano WL. The PyMOL Molecular Graphics System. CCP4 Newsletter on protein crystallography. 2002;40(1).

